# Exploring Proteomic Differences in PBMCs for Sex-Specific Insights into Alzheimer’s Disease

**DOI:** 10.1101/2025.10.31.685960

**Authors:** Takako Ishida-Takaku, Colin K. Combs

## Abstract

Peripheral blood mononuclear cells (PBMCs) offer a minimally invasive window into systemic biology and immune dysregulation in Alzheimer’s disease (AD). We performed quantitative proteomic profiling of PBMCs from male and female AD patients and controls to assess sex differences. AD was associated with proteomic remodeling, with complement activation, coagulation, and neuronal signaling enriched in males, whereas females showed increased steroid hormone secretion, lipid metabolism, and acute-phase response with reduced translation and DNA maintenance. Despite distinct patterns, both sexes exhibited immune and hemostatic activation, underscoring shared systemic mechanisms and the need for sex-specific biomarkers and therapeutic strategies in AD.

## Description

AD is primarily a central nervous system disorder, and systemic immune alterations contribute substantially to its pathogenesis (Bettcher et al., 2021; Heneka et al., 2015; Van Eldik et al., 2016). PBMCs, which include lymphocytes and monocytes, reflect immune status and inflammatory signaling in the body and can mirror aspects of brain pathology. Studies have shown that PBMCs from AD patients exhibit changes in gene expression (Fiala et al., 2007), cytokine production (Pellicanò et al., 2010), oxidative stress responses (Mecocci et al., 2002), and proteomic profiles, making them a minimally invasive surrogate for detecting peripheral and potentially central alterations. In this study, we performed proteomic analysis of PBMCs from male and female AD patients and from age-matched male and female controls with incidental age-related pathology (n = 6 to 8) to explore sex specific peripheral proteomic alterations in AD. Differential expression and pathway analyses were performed using iDEP v2.01 (Ge, 2021; Wu et al., 2025), an integrated web-based platform for omics data exploration. Differential expression analysis with lenient thresholds (raw p value < 0.1, Fold Change ≥ 1.5) identified 53 upregulated and 106 downregulated proteins in AD males compared with male controls (Figure 1A), whereas AD females exhibited 191 upregulated and 369 downregulated proteins compared with female controls (Figure 1B). Direct comparison of AD males and females identified 103 proteins upregulated in males and 56 upregulated in females (Figure 1C).

**Figure 1.**
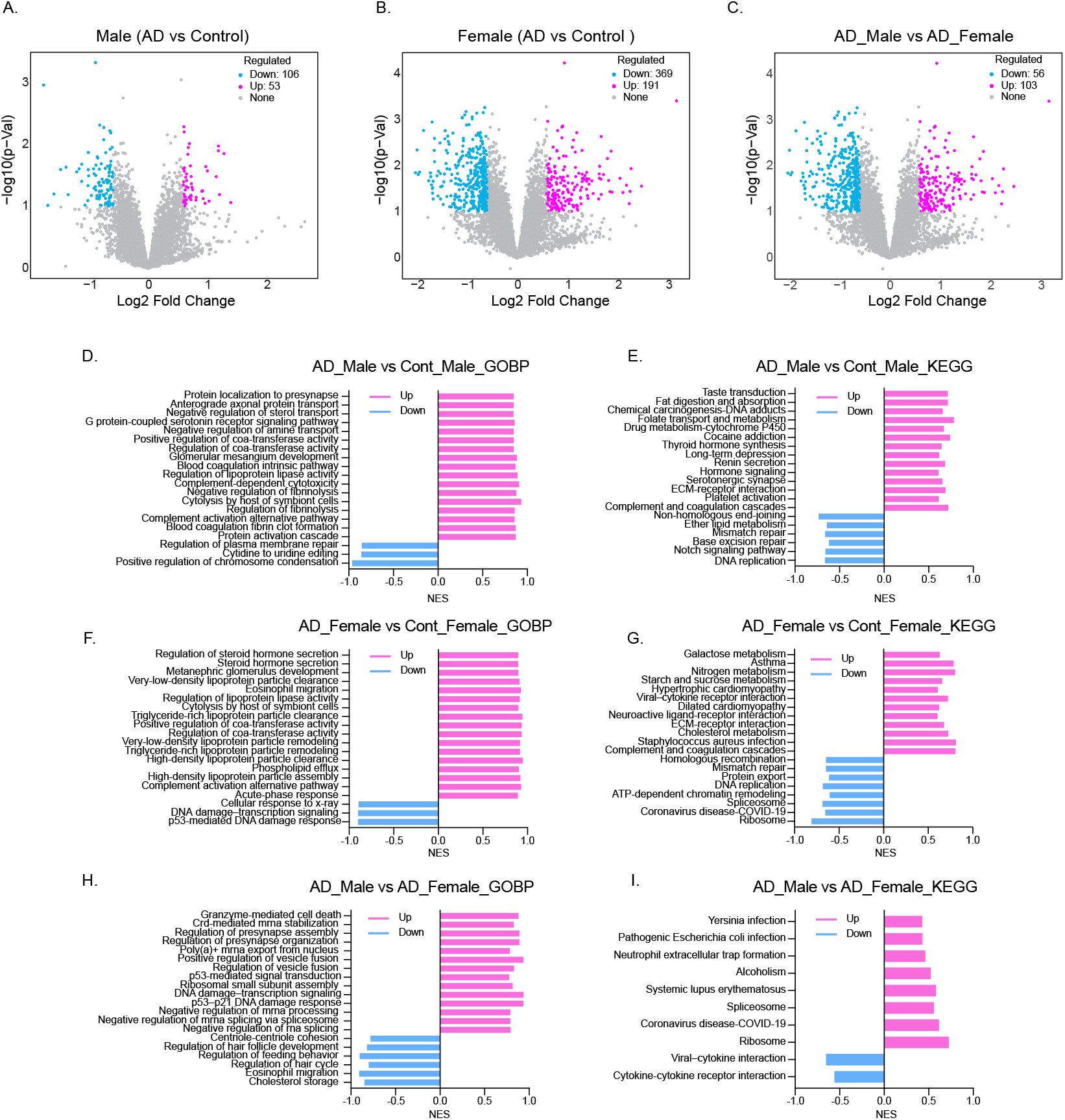
Differential proteomic analysis and pathway enrichment in PBMCs from Alzheimer’s disease (AD) patients and controls. (A–C) Volcano plots showing log2 fold change versus –log10 low p-value for proteins detected in comparisons of AD male vs. control male (A), AD female vs. control female (B), and AD male vs. AD female (C). Numbers of upregulated, downregulated, and unchanged proteins are indicated. (D–F) Gene Ontology Biological Process (GOBP) enrichment plots for significantly regulated proteins in the same comparisons. (G–I) KEGG pathway enrichment plots for the corresponding comparisons. Analyses were performed using the iDEP 2.1 web-based platform. Significance of enrichment wasdefined as FDR < 0.1 based on adjusted p-values, and normalized enrichment scores (NES) were used to rank pathways.

Pathway analyses Gene Set Enrichment Analysis (GSEA)(Alhamdoosh et al., 2017), in which normalized enrichment scores (NES) are shown and pathways with FDR < 0.1 were considered statistically significant, revealed both shared and sex-specific biological signatures in AD. In AD males, Gene Ontology Biological Process (GO BP) analysis indicated enrichment of pathways involved in neural signaling (e.g., presynaptic localization, axonal protein transport, serotonin receptor signaling), as well as coagulation and complement activation, suggesting AD-associated alterations in both neuronal and immune-related processes (Figure 1D). KEGG analysis further identified hormone signaling pathways as upregulated, while DNA replication and repair pathways were suppressed, in addition to alterations in neuronal and immune pathways (Figure 1E).

In AD females, by contrast, the most pronounced changes included steroid hormone secretion, lipoprotein particle clearance, acute-phase response, and complement activation (Figure 1F). KEGG analysis further highlighted upregulation of steroid hormone–related signaling, lipid and carbohydrate metabolism (galactose, nitrogen, starch, and sucrose), cholesterol metabolism, and complement and coagulation cascades (Figure 1G). Despite these sex-specific emphases, males and females exhibited enrichment of complement activation, coagulation cascades, and extracellular matrix (ECM) receptor interaction pathways. Notably, proteins involved in DNA replication, repair, ribosome, and spliceosome pathways were broadly downregulated in female AD, whereas male AD showed less pronounced suppression of these processes.

A direct comparison of AD males versus AD females revealed divergent biological programs. GO BP terms enriched in males included granzyme-mediated cell death, p53-mediated signaling, vesicle fusion and presynapse organization, and ribosomal subunit assembly, whereas females showed enrichment of cholesterol storage, eosinophil migration, and developmental/behavioral regulation (Figure 1H). KEGG pathways upregulated in males included pathogen-related infections, neutrophil extracellular trap formation, systemic lupus erythematosus, spliceosome, and ribosome, while females were enriched for cytokine and cytokine receptor interaction and viral protein interaction with cytokine and cytokine receptor (Figure 1I). These results show sex dependent differences in PBMC biology, which may influence how peripheral changes are linked to AD pathology.

Our findings show that PBMCs from AD patients show sex-dependent changes in protein expression, with some alterations shared between males and females and others unique to each sex. In males, AD was associated with enrichment of neuronal and immune-related pathways and modest suppression of DNA replication and repair. This pattern suggests that male PBMCs are biased toward immune and hemostatic activation and immunosenescence, consistent with vascular dysfunction and complement dysregulation reported in AD (Haage & De Jager, 2022; Mehta & Mehta, 2023; Rodrigues et al., 2021). In contrast, females showed prominent activation of steroid hormone secretion, lipid metabolism, and complement pathways, accompanied by marked downregulation of DNA replication, repair, and ribosomal functions. These results suggest that female PBMCs display increased metabolic and inflammatory responsiveness but reduced translational and genomic maintenance capacity, reflecting systemic stress and proteostasis imbalance. Complement activation, coagulation, and ECM–receptor interaction pathways shared between males and females represent common systemic features of AD (Haytural et al., 2021; Kodam et al., 2023), reflecting coordinated immune, hemostatic, and structural remodeling processes. Direct comparison between the sexes revealed that AD males preferentially upregulate cytotoxic and translational immune programs, including granzyme-mediated cell death and ribosomal and spliceosome activity, while AD females show cytokine signaling and lipid-related processes. This alteration may be driven by hormonal, metabolic, and immune regulatory mechanisms, and could underlie differences in AD susceptibility, progression, and therapeutic responses between men and women. Taken together, the data show potential sex dependence in peripheral AD biology, and our findings suggest that a single biomarker panel may be inadequate and that early diagnostic biomarkers may need development on a sex specific basis to capture the dominant biology in each group. Larger and longitudinal cohorts, cell type resolved proteomics, and analyses that combine PBMC with brain and plasma datasets will test generalizability and refine sex informed strategies for diagnosis and treatment.

## Methods

### Human PBMCs

Human peripheral blood mononuclear cells (PBMCs) were obtained from Banner Health. PBMCs from Alzheimer’s disease (AD) patients were collected from individuals aged 52 years or older. Age-matched control samples were obtained from individuals aged 71 years or older.

### Quantitative Proteomics

Proteins were reduced, alkylated, and subjected to chloroform/methanol extraction before enzymatic digestion with sequencing-grade modified trypsin (Promega). The resulting peptides were separated on a reverse-phase Ion-Opticks-TS analytical column (25 cm × 75 µm, 1.7 µm C18 resin) coupled to an EASY-Spray source maintained at 60°C. Samples were loaded onto a PepMap Neo trap column (300 µm × 5 mm) using a Vanquish Neo UHPLC system (Thermo Scientific) at 11°C before injection. Peptides were resolved over a 35-min gradient at 0.35 µL/min, starting from 98% buffer A (0.1% formic acid, 0.5% acetonitrile in water) and 2% buffer B (80% acetonitrile, 20% water, 0.1% formic acid), with stepwise increases to 56:44 at 27.1 min, followed by column washing and re-equilibration. Eluted peptides were ionized at 2.5 kV and analyzed on an Orbitrap Astral mass spectrometer (Thermo Scientific) operated in DIA mode. MS1 spectra were collected across 380–980 Th at 240,000 resolutions with a normalized AGC target of 200% and a maximum injection time of 3 ms. DIA scans consisted of 199 windows (3 Th each) with 25% HCD collision energy, normalized AGC target of 100%, and maximum injection time of 3 ms. MS2 spectra were acquired from 150–2000 Th with the RF lens set at 40%.

### Data Processing and Statistical Analysis

Spectral data were processed in Spectronaut (Biognosys v19.5) against the UniProt *Homo sapiens* reference proteome (UP000005640, release 2025) using the directDIA workflow. A 1% precursor and protein-level q-value cutoff was applied, with decoys generated for FDR control. Protein inference was performed with the IDPicker algorithm, using MS2-level quantification and median peptide/precursor intensities (Searle et al., 2018). Protein MS2 intensity values were assessed for quality using ProteiNorm (Graw et al., 2020). The data were normalized using VSN (Huber et al., 2002) and analyzed using proteoDA to perform statistical analysis using Linear Models for Microarray Data (limma package (Ritchie et al., 2015)) with empirical Bayes (eBayes) smoothing to the standard errors (Chawade et al., 2014; Thurman et al., 2023).

## Acknowledgements

We are grateful to Angela Floden for assistance with sample management for this proteomic analysis. We also acknowledge the IDeA National Resource for Quantitative Proteomics, supported by NIH/NIGMS grant R24GM137786, for expert support in mass spectrometry-based proteomics analysis.

## Author Contributions

Takako Ishida-Takaku: Data analysis, Visualization, Writing – original draft and editing, Colin K. Combs: Conceptualization, Funding acquisition, Supervision, Writing - review and editing

